# CGMTSA: An R package for continuous glucose monitoring time series data analysis

**DOI:** 10.1101/2020.07.06.174748

**Authors:** Jian Shao, Tao Xu, Kaixin Zhou

## Abstract

The R package CGMTSA was developed to facilitate investigations that examine the continuous glucose monitoring (CGM) data as a time series. Accordingly, novel time series functions were introduced to: 1) enable more accurate missing data imputation and outlier identification; 2) calculate recommended CGM metrics as well as key time series parameters; 3) plot interactive and 3D graphs that allow direct visualizations of temporal CGM data and time series model optimization. The software was designed to accommodate all popular CGM devices and support all common data processing steps.

## 1 Introduction

Continuous glucose monitoring (CGM) is an effective tool to measure glucose concentration in the interstitial fluid at a relatively short gap of 5-15 minutes over a few days. Using CGM in patients with diabetes could improve glycaemic control owing to the continuous monitoring (Battelino et al., 2019). Recently, a few software tools had been developed to facilitate more consistent estimation of CGM data metrics (Zhang et al., 2018; Millard et al., 2020) and demonstrating various glucose fluctuation metrics associated with diabetes complications (Ceriello et al., 2019).

However, CGM by nature produces time series data which could be decomposed into three components of trend, periodic and residuals originated from random life events. Time series analysis of this type of data has been fruitful in other fields such as stock price prediction. It is expected comprehensive mining of CGM time series data could lead to breakthrough in accurate risk prediction of glycaemic event and long-term complications. Here we developed a software for Continuous Glucose Monitoring Time Series Data Analysis (CGMTSA) to provide essential functions that facilitate time series analyzes of CGM data.

## 2 Features

CGMTSA provides three groups of functions for quality control, metrics calculation and data visualization. Apart from common functions as provided by other software, we also implemented methods to bring out the time series features of CGM data.

### 2.1 Quality control

CGMTSA can read original data from devices made by all major suppliers which take different formats such as Abbott FreeStyle Libre, Dexcom G6 and Medtronic Ipro2. To tackle the challenge of missingness, three imputation methods were provided, including linear regression, autoregressive integrated moving average model (ARMA) and seasonally decomposed missing value imputation model (SEADEC). ARMA and SEADEC are both time series models that have been used to make predictions with and without the periodic component respectively. Similarly, ARMA was also used in CGMTSA to identify CGM data outliers, which often originate from mechanical pressure on the sensor or patient motion that could significantly bias data interpretation (Facchinetti et al., 2016). The processed CGM data points will be labelled with these quality control flags.

### 2.2 Metrics calculation

CGMTSA outputs most commonly used glycaemic metrics. The measures of blood glucose variability presented include standard deviation (SD), coefficient of variation (CV), mean of daily difference (MODD), time in range (TIR) and mean amplitude of glycemic excursion (MAGE) which are recommended by international consensus on the use of CGM (Danne et al., 2017). CGMTSA also calculates LBGI and HBGI that are recommended to estimate the risk of hypoglycemia and hyperglycemia (Kovatchev et al., 1997). For key time series parameters of autocorrelation coefficients (ACF) and partial autocorrelation coefficients (PACF), CGMTSA outputs the first five sequential coefficients in ACF and PACF. By estimating correlations between current values of a time series and its lagged values, coefficients of ACF and PACF could be used to optimize the performance of time series regression models.

### 2.3 Data visualization

CGMTSA constructs an interactive three-dimensional plot of date, time and glucose level (Fig. 1B). This will allow investigators direct visualizations of the day-to-day change in glucose levels at various time points, which could hardly be achieved in traditional 2D plots. To help validate the assumption of stationarity in a time series data set, CGMTSA calculates the statistics of augmented Dickey–Fuller test and plots autocorrelation coefficients of ACF and PACF (Fig. 1C).

**Fig. 1.**
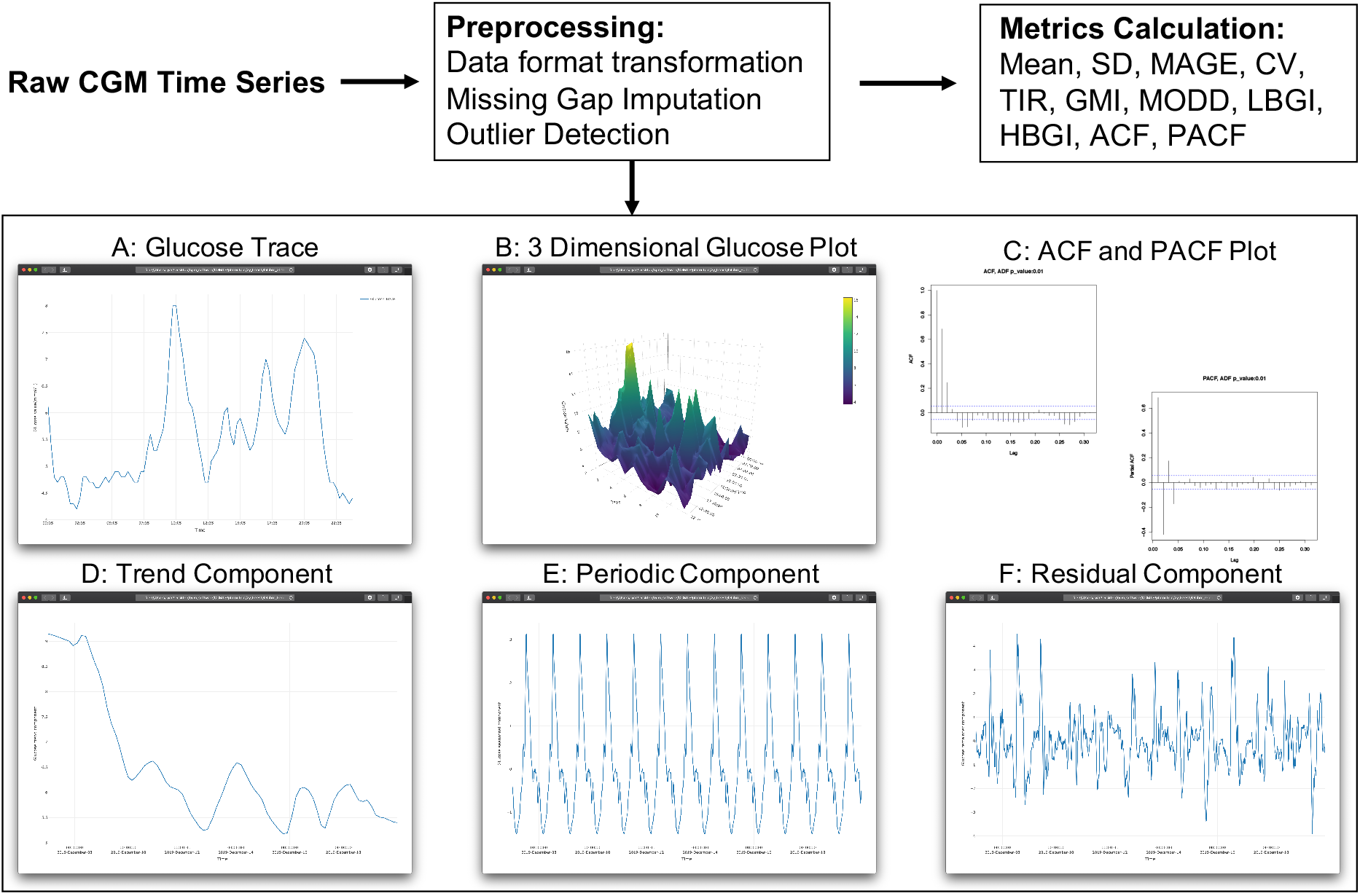
Workflow of continuous glucose monitoring time series data analysis. The detected outliers will be labeled in different colors in glucose trace plot.

To disentangle the complexity of CGM data, CGMTSA uses additive decomposition to plot the trend, periodicity and residual components as interactive plots. The trend measures the change of blood glucose over sliding time windows (Fig. 1D). The periodic component averages the detrended glucose values at certain time point across days to reflect the periodicity of glucose fluctuation (Fig. 1E). After removing the trend and periodic background, the residual component plot could highlight random glycaemic events such as those induced by medical or lifestyle interventions (Fig. 1F).

## 3 Results

To demonstrate a typical use case for CGMTSA, we utilized one example of CGM data. This example is from Abbott Freestyle Libre contains 14 days glucose data. The original format and preprocessed format of data are presented in Fig.2. The sglucose column is glucose values from CGM sensor. The bglucose column is self-monitoring blood glucose value. The imglucose column is imputed glucose values from sglucose. The outlier column are glucose values that are considered outlier by time series outlier detection algorithm.

**Fig. 2.**
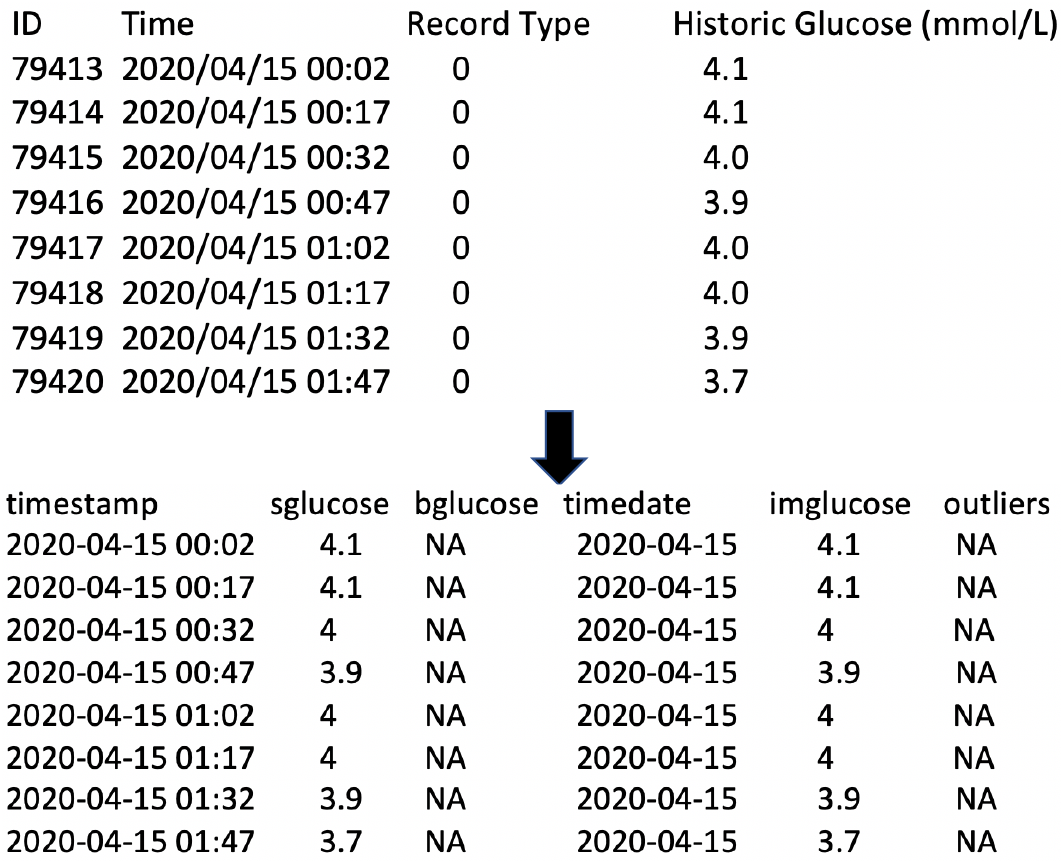
Data format transformation. The original format of Abbott Freestyle Libre includes time, record type and historic glucose. The preprocessed format includes timestamp, sensor glucose (sglucose), blood glucose (bglucose), time date, imputed glucose (imglucose) and outliers.

**Fig. 3.**
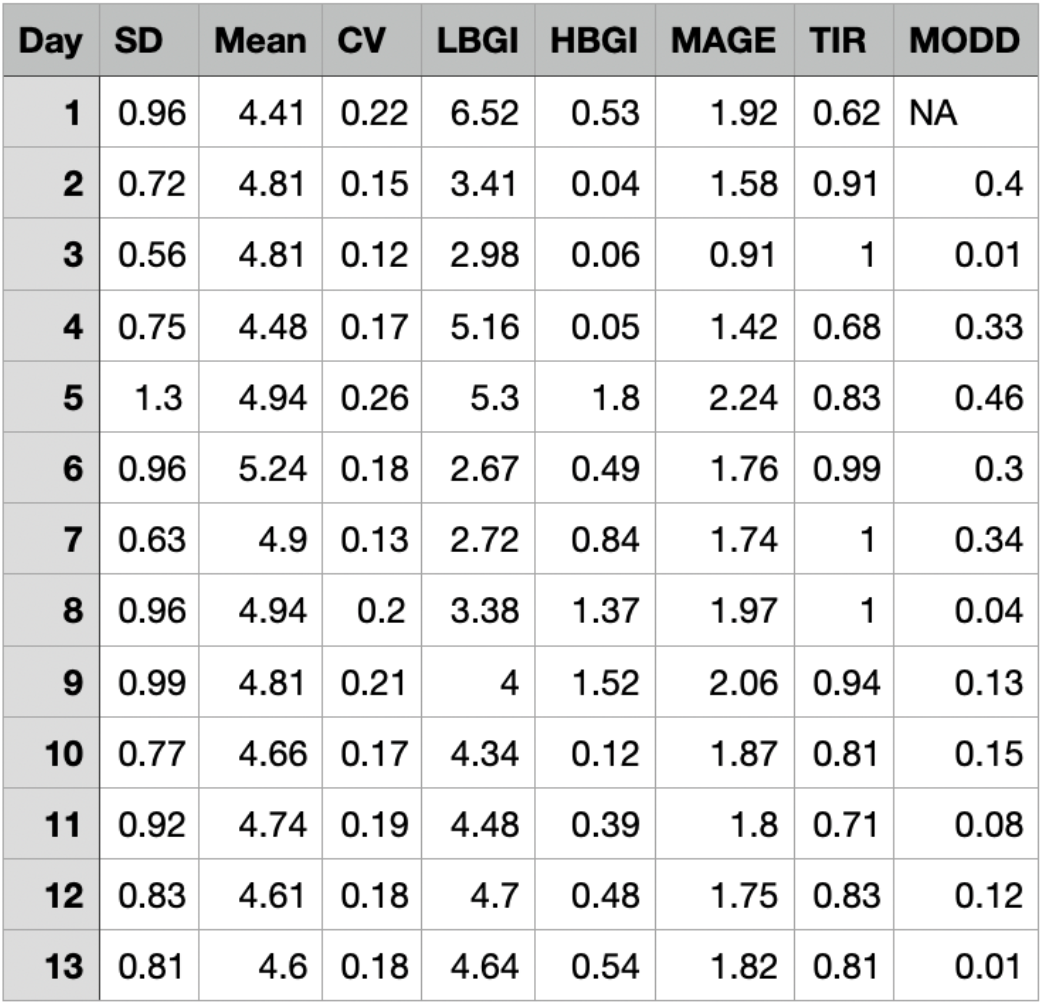
Glucose metrics. The metrics includes standard deviation (SD), mean of glucose, coefficient of variation (CV), low blood glucose index, high blood glucose index, mean of daily difference (MODD), time in range (TIR) and mean amplitude of glycemic excursion (MAGE).

After quality control, we use CGMTSA to estimate metrics of glucose fluctuation. The metrics includes standard deviation (SD), mean of glucose, coefficient of variation (CV), low blood glucose index, high blood glucose index, mean of daily difference (MODD), time in range (TIR) and mean amplitude of glycemic excursion (MAGE) which are recommended by international consensus on the use of CGM (Fig. 2). These metrics are estimated by day, the summary metrics of all days also are estimated. CGMTSA outputs the first five sequential coefficients in ACF and PACF (Supplemental Table 1). ACF and PACF coefficients can be used in construction of the autoregressive model.

The glucose data of many days are overlapped in a 2-dimensional plot in traditional glucose plot. In 2-dimensional plot, the day-to-day change in glucose levels at various time points could hardly be achieved. To overcome this gap, CGMTSA constructed an interactive three-dimensional plot of date, time and glucose level (Fig.4). Fig.4B shows two different angles of 3-dimensional plot, the peak of waves of this example are closed to each other which indicate the day-to-day change is small.

**Fig. 4.**
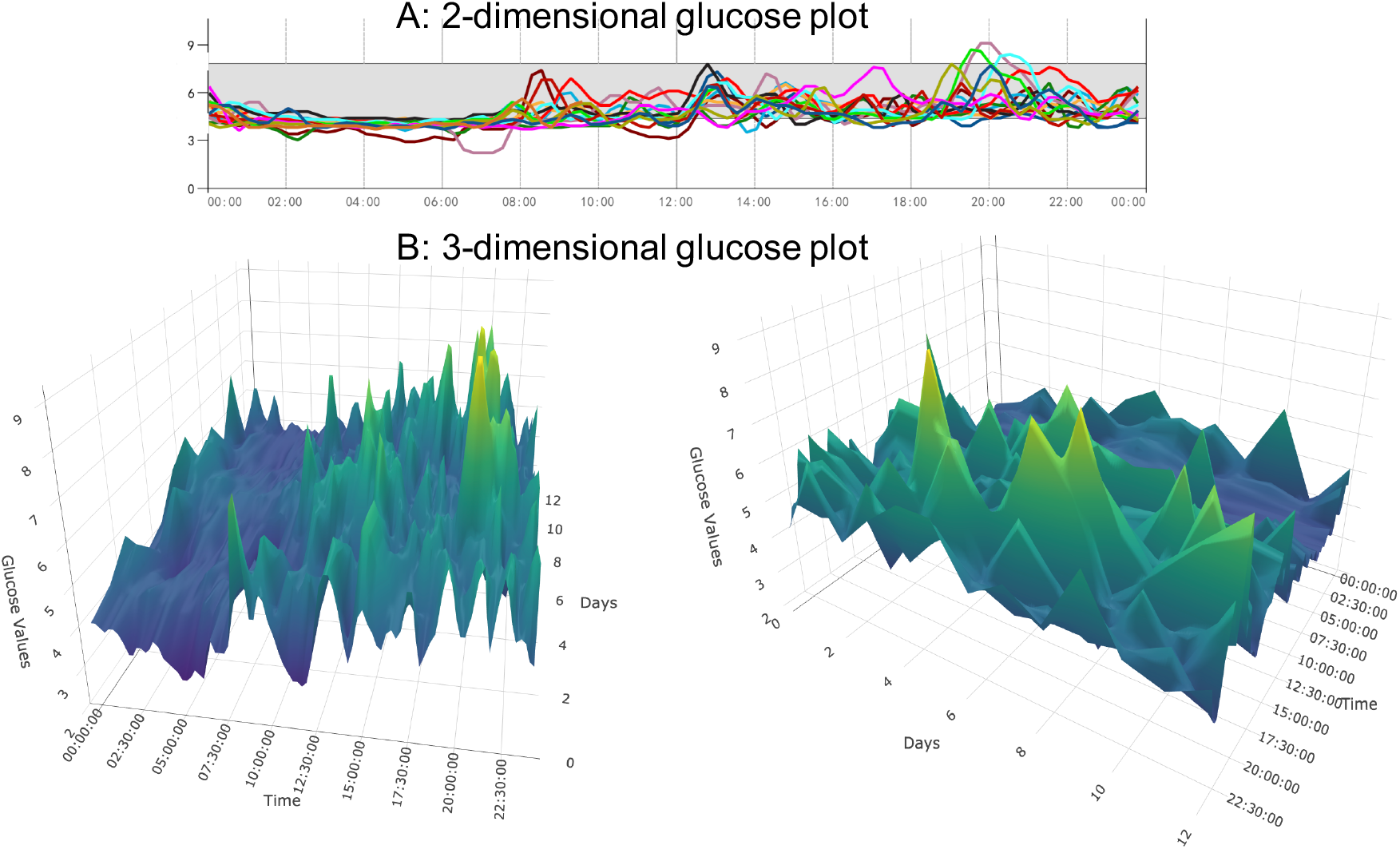
Three-dimensional glucose plot of preprocessed 14 days glucose data. Fig. 4A shows the traditional 2-dimensional plot. Fig. 4B shows the interactive 3-dimensional plot.

CGMTSA uses time series additive decomposition algorithm to plot the trend, periodicity and residual components as interactive plots. Fig.5 shows the result of decomposition from preprocessed CGM data. Trend component shows glucose of this example reached the highest in the middle of 14 days and decreased after this point (Fig. 5A). Periodic component shows that mean of glucose values are lower at six than other times, mean of glucose values are higher at eight than other times (Fig. 5B). The residual component reflects random glycaemic events such as those induced by medical or lifestyle interventions (Fig. 5C). CGMTSA also outputs the ACF and PACF plot which can help validate the assumption of stationarity in a time series data set (Supplemental Figure 5,6).

**Fig. 5.**
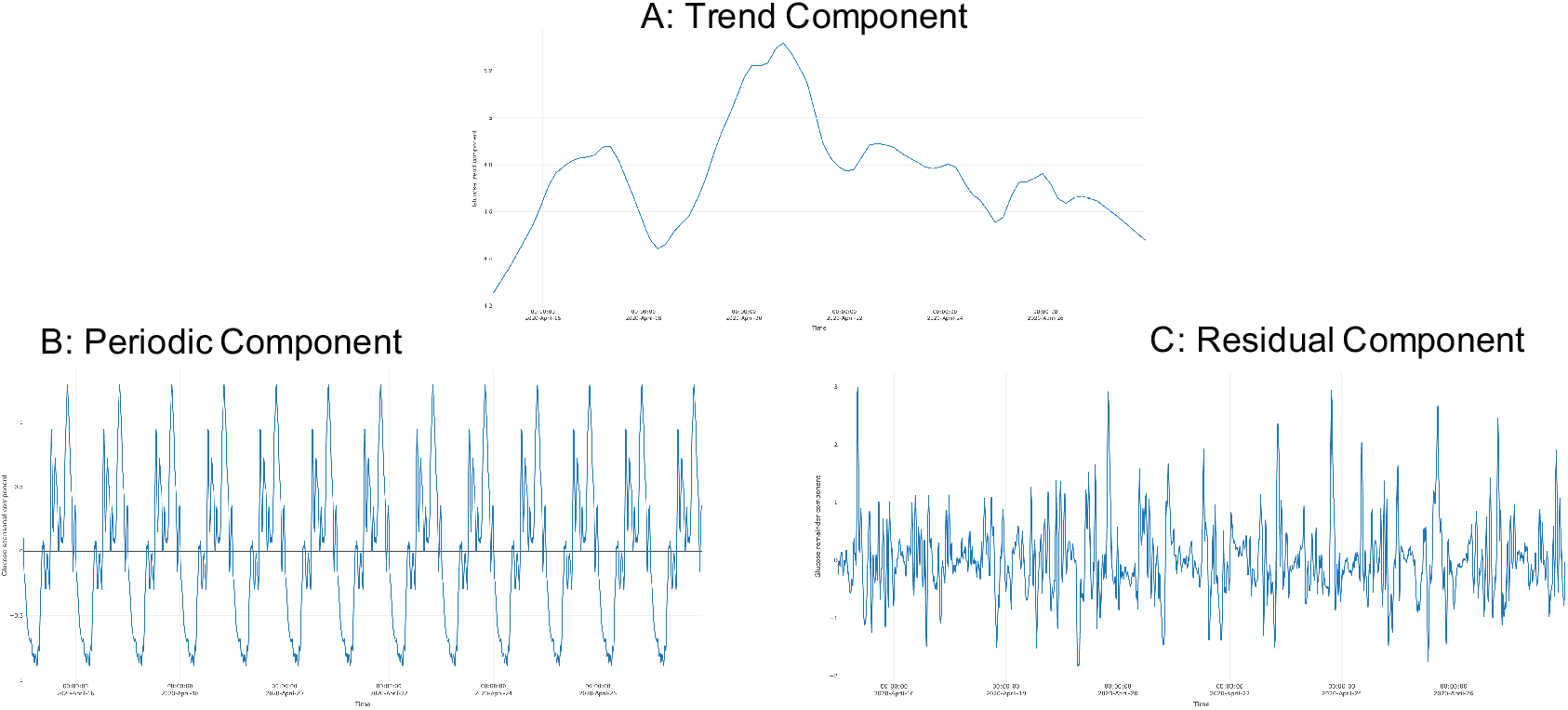
Results of CGM data decomposition.

## 4 Conclusion

We presented an open-source R package CGMTSA for more comprehensive analyses of time series data from all common CGM devices. With a focus on dissecting the temporal characteristics embedded in CGM data, specific functions for quality control, metrics calculation and visualization were introduced to pave the way for novel downstream clinical investigations. Compared to other CGM software that focuses on amplitude and fluctuations of glucose, such time series analyses can derive more reliable data sets which would enable more robust statistical modelling and accurate risk predictions.

## Availability and implementation

The CGMTS package, additional documentation and example datasets are available at https://github.com/RyanJ-Shao/CGMTSA. The software is open-source and released under the MIT License.

## Notes

https://github.com/RyanJ-Shao/CGMTSA

### Competing Interest Statement

The authors have declared no competing interest.

