## Supplementary figures and images for "CGMTSA: An R package for continuous glucose monitoring time series data analysis"

### Supplemental Figure 5

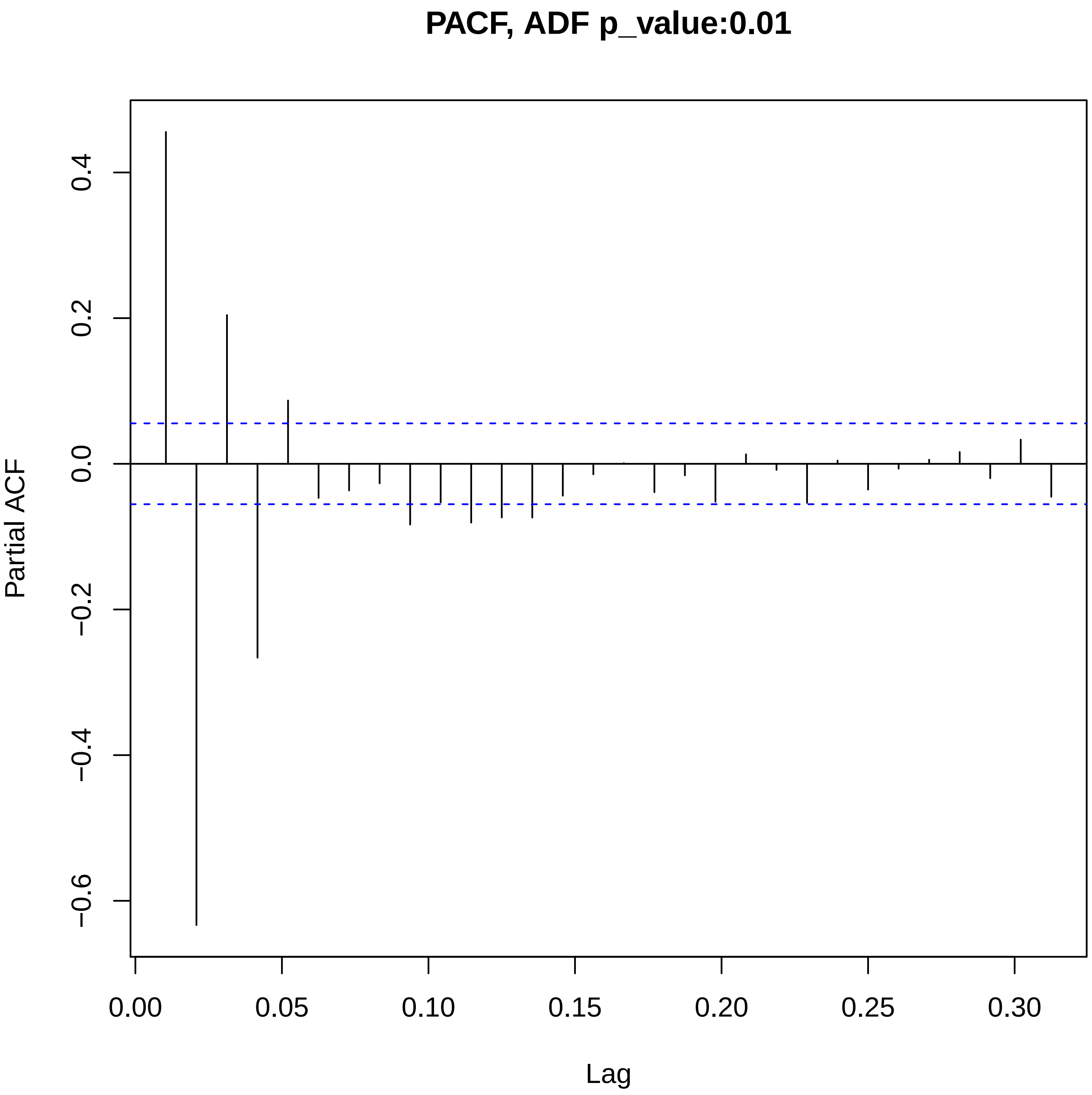

### Supplemental Figure 6

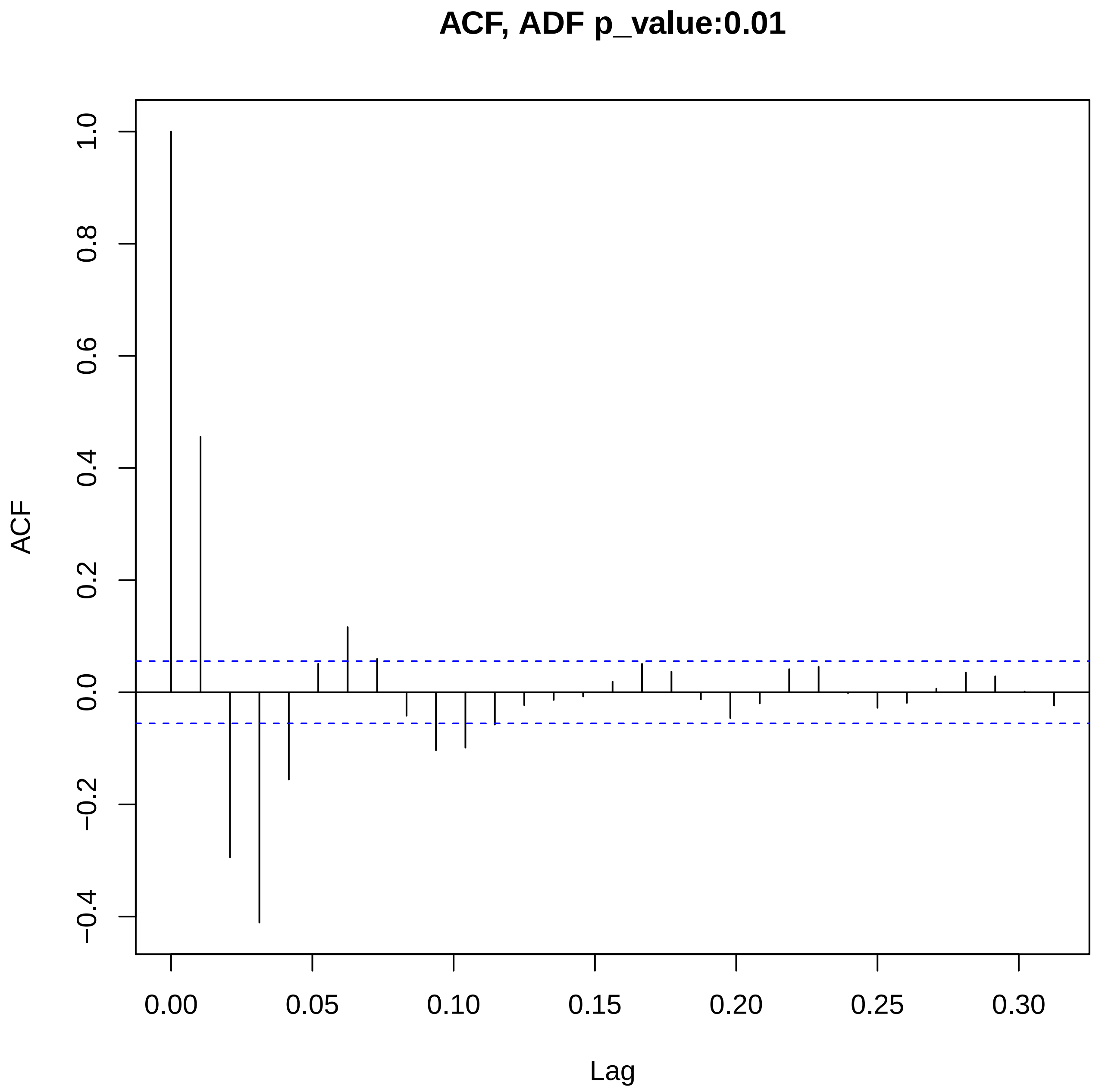
